# RB loss sensitizes cells to replication-associated DNA damage by PARP inhibition

**DOI:** 10.1101/2023.03.25.532215

**Authors:** L Gregory Zamalloa, Margaret M Pruitt, Nicole M Hermance, Himabindu Gali, Rachel L Flynn, Amity L Manning

**Author notes:** materials requests and correspondence should be addressed to: Amity L Manning, 60 Prescott St Rm 4002, Worcester MA 01605, USA.

## Abstract

The retinoblastoma tumor suppressor protein (RB) interacts physically and functionally with a number of epigenetic modifying enzymes to control transcriptional regulation, respond to replication stress, promote DNA damage response and repair pathways, and regulate genome stability. To better understand how disruption of RB function impacts epigenetic regulation of genome stability and determine whether such changes may represent exploitable weaknesses of RB-deficient cancer cells, we performed an imaging-based screen to identify epigenetic inhibitors that promote DNA damage and compromise viability of RB-deficient cells. We found that loss of RB alone leads to high levels of replication-dependent poly-ADP ribosylation (PARylation) and that preventing PARylation through inhibition of PARP enzymes enables RB-deficient cells to progress to mitosis with unresolved replication stress and under-replicated DNA. These defects contribute to high levels of DNA damage, decreased proliferation, and compromised cell viability. We demonstrate this sensitivity is conserved across a panel of inhibitors that target both PARP1 and PARP2 and can be suppressed by re-expression of the RB protein. Together, these data indicate that inhibitors of PARP1 and PARP2 may be clinically relevant for RB-deficient cancers.

## Introduction

The retinoblastoma tumor suppressor (RB), which is lost or functionally inactivated in the majority of cancers (Burkhart and Sage 2008), is best known for its role as a negative regulator of E2F-dependent transcription (Dyson 1998; Nevins 2001). As RB becomes increasingly phosphorylated during G1 progression, its inhibition of E2F is abrogated, allowing for expression of key cell cycle genes, progression through the restriction point, and S-phase entry (Goodrich et al. 1991). However, the cellular function of RB is now appreciated to be much more extensive than E2F regulation (Velez-Cruz and Johnson 2017; Dick et al. 2018) and proteomics analyses indicate that RB interacts with >300 proteins, nearly 20% of which are histones or epigenetic modifiers of histones (Sanidas et al. 2019).

Interactions with epigenetic regulators are believed to allow RB to orchestrate chromatin accessibility and transcription status across the cell cycle (Gonzalo and Blasco 2005; Guzman et al. 2020). Reported RB interactors of this nature include histone acetylases (Manickavinayaham et al. 2019), deacetylases (Luo et al. 1998; Wang et al. 2019; Zhou et al. 2021), methylases, and demethylases (Nielsen et al. 2001; Vandel et al. 2001; Gonzalo et al. 2005; Blais et al. 2007; Chau et al. 2008; Ishak et al. 2016). Disruption of RB-dependent regulation of epigenetic factors compromises transcriptional regulation, induces replication stress, impairs DNA damage response and repair pathways, and corrupts genome stability (Manickavinayaham et al. 2020). These data suggest epigenetic dysregulation may be a critical and exploitable feature of RB-deficient cells.

Indeed, recent reports demonstrate that RB-deficient cells are exquisitely sensitive to inhibition of epigenetic modulators, including the key mitotic kinases Aurora A and Aurora B (Gong et al. 2019; Oser et al. 2019; Lyu et al. 2020; Yang et al. 2022). These kinases phosphorylate key regulators of centromere and spindle structure and function, and their inhibition synergizes with RB-dependent defects in centromere regulation and chromosome segregation (Iovino et al. 2006; Amato et al. 2009; Manning et al. 2010; Schvartzman et al. 2011; Manning et al. 2014). Other studies have described sensitivity of RB-deficient cells to pharmacological inhibition of polo-like kinase 1 (PLK1) function such that RB-proficient cells arrest in response to PLK1 inhibition while those deficient for RB continue to proliferate, accumulate high aneuploidy, and ultimately undergo cell death (Witkiewicz et al. 2018).

Prior studies have also implicated RB loss in conferring sensitivity to inhibition of poly ADP ribose polymerase (PARP) enzymes (Velez-Cruz et al. 2016; Jiang et al. 2020; Zoumpoulidou et al. 2021), yet the mechanistic explanation for this relationship remains unclear. Here we identify PARP1/2 inhibitors in a screen for epigenetic modulators that exploit DNA damage phenotypes in cells lacking RB. We utilize isogeneic cell lines with and without RB to show that RB deficiency leads to high levels of replication-dependent PARylation. PAR, an epigenetic mark placed by PARP enzymes, marks sites of Okazaki fragment processing and replication stress. This modification functions to activate the DNA damage response pathways and stall replication until the stress is resolved (Ame et al. 2004; Sugimura et al. 2008; Bryant et al. 2009; Hanzlikova et al. 2018; Vaitsiankova et al. 2022). We find that inhibition of PARP activity permits acceleration of replication fork progression in a manner that is ignorant of RB-dependent DNA lesions. Progression into G2/M in the presence of unresolved DNA damage contributes to genomic instability and compromised cell viability. Restoration of RB expression in the isogenic cell line is sufficient to protect cells from replication stress associated with PARP inhibition.

## Results

### RB-deficient cells are sensitive to PARP1 inhibition

Previous studies have found that RB plays a role in replication fork progression and homologous recombination (Marshall et al. 2019) and that RB-deficient cells are sensitive to DNA damaging agents that generate double strand breaks (Velez-Cruz et al. 2016; Aubry et al. 2020; Jiang et al. 2020). Regulation of chromatin structure is critical to the repair of DNA damage and modulation of nucleosome positioning at sites of breaks is shown to be limiting for repair (Hauer and Gasser 2017). Therefore, we hypothesized that defects in replication and homologous recombination that result from RB loss may render cells sensitive to epigenetic perturbations of chromatin structure. To test this possibility, we performed a targeted small molecule screen to assess measures of genome stability and viability following exposure to individual inhibitors from a panel of 96 small molecule modulators of epigenetic regulation. We designed a small molecule library to target 20+ categories of epigenetic modulators, including HDMs, HDACs, DNMTs, kinases and PARPs (Supplemental Table S1), that have been previously implicated in the maintenance of chromatin structure and genome stability (Putiri and Robertson 2011; Sultanov et al. 2017; Karakaidos et al. 2020).

Using an hTERT RPE-1 (RPE) cell line in which RB can be depleted through doxycycline induced expression of an RB-targeting shRNA (Manning et al. 2014), cells were first depleted of RB for 48 hours. Populations of cells with and without RB depletion were then exposed to individual epigenetic modulators for 48 hours and assessed for early signs of sensitivity, including acquisition of DNA damage and reduced cell number. Seventeen inhibitors that caused significant death of control cells (defined as 80% or greater reduction in cell number after 2 cell cycles) were not evaluated further. To assess DNA damage following exposure to the remaining 79 inhibitors, cells were immunostained for γH2AX, a phosphorylated histone mark and early indicator of DNA double strand breaks. Using Nikon elements software, individual nuclei were identified by thresholding in the DAPI channel and sum intensity of γH2AX staining was measured (Supplemental Fig. S1A) to assess levels of DNA damage. Individual cells were considered to have enhanced DNA damage if nuclear γH2AX staining was more than 2-fold the average γH2AX intensity measured in control cells from the same screening plate. Relative fold change in the fraction of damaged cells was calculated to identify modulators that cooperate with RB loss to promote DNA damage. Fourteen inhibitors differentially induced a 2-fold or greater increase in DNA damage in cells depleted of RB with a robust Z-score of 3.0 or greater across biological triplicates (Fig. 1A,B).

**Figure 1.**
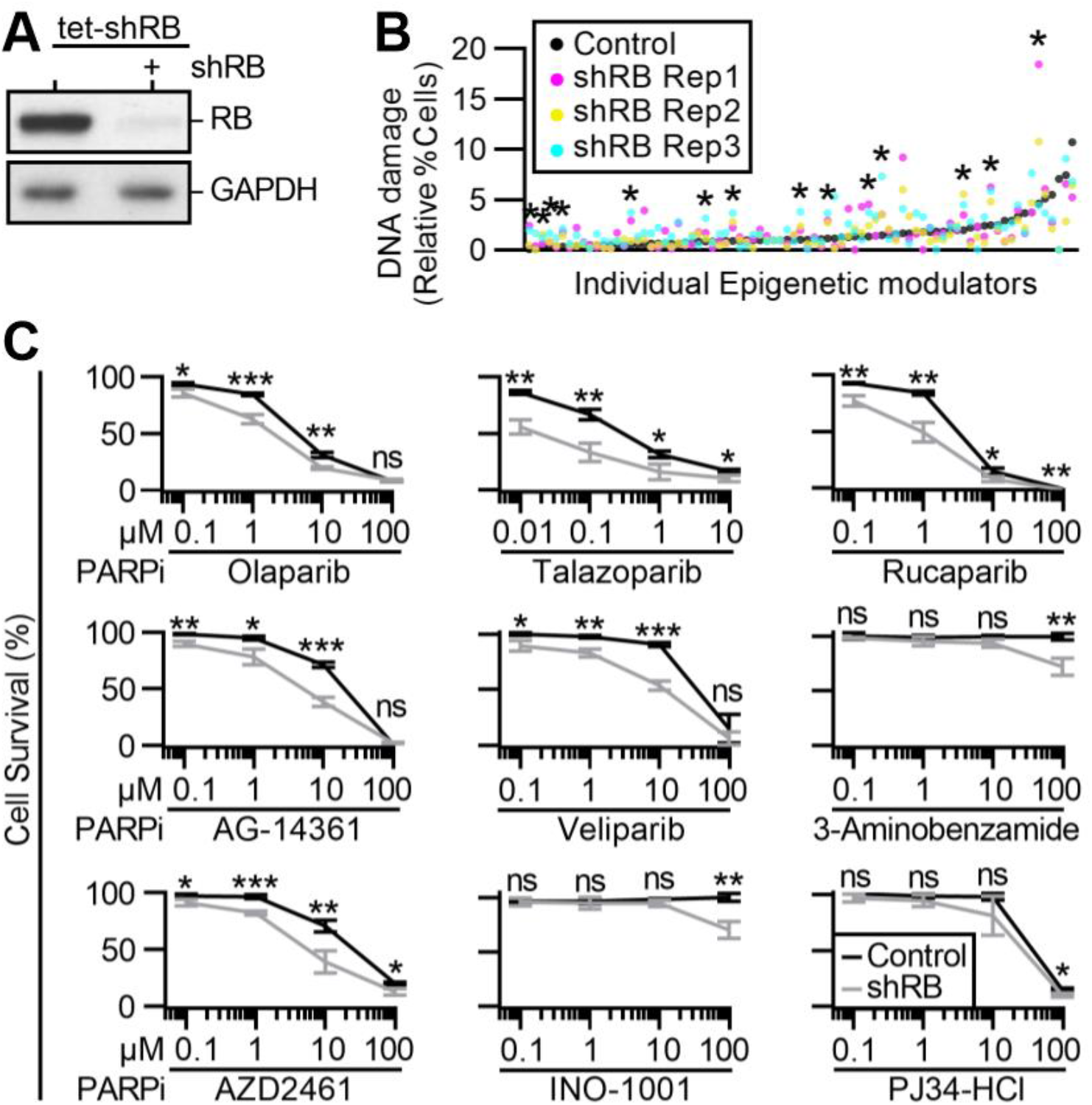
Identification of epigenetic sensitives of RB-deficient cells. *(A)* Western blot analysis of cells with and without 96h induced expression of an RB-targeting shRNA (tet-shRB). *(B)* Fraction of control (black) and shRB (gray) cells with DNA damage following 48h exposure to individual epigenetic modulators. * Indicates differential increase in DNA damage in RB-deficient cells with robust Z score > 3 *(C)* Relative cell survival of control and shRB cells, as indicated by metabolic color conversion of PrestoBlue reagent following incubation with the indicated PARP inhibitors and concentrations. All experiments were performed in biological triplicate and statistics performed between biological replicates. (*) p < 0.05; (**) p < 0.01; (***) p < 0.001; (ns) non-significant p > 0.05.

Recent reports have indicated that RB-deficient cells are exquisitely sensitive to inhibition of Aurora kinase, EZH2 methyltransferase, and BRD4 inhibition (Ishak et al. 2016; Gong et al. 2019; Oser et al. 2019; Zhang et al. 2021). Consistent with these results, our screen identifies inhibitors of Aurora kinase (via JNJ-7706621 and CYC116), inhibitors of EZH2 (via EPZ-6438 and 3-Deazaneplanocin A (DZNeP)), and inhibitors of BRD4 (via PFI-1 and Bromosporine) that each differentially enhance DNA damage levels in RB-depleted cells. Additional small molecules that scored as hits in our screen were inhibitors of HDAC (RGFP966, Rocilinostat, and Resminostat), JAK2 (Ruxolitinib, AZD1480, and Gandotinib), and PARP (Olaparib and Talazoparib). To further characterize these hits, we next assessed cell viability following exposure to a concentration range of each drug (Supplemental Fig. S1B). Of the fourteen inhibitors assessed for impact on viability, only Olaparib and Talazoparib, both PARP1/2 inhibitors, differentially compromised viability of RB-deficient RPE cells in the 48-96h time course of this experiment (Supplemental Table S2).

Seven additional PARP inhibitors were represented in the screening library but did not meet the criteria described above to be considered hits on the screen. We therefore sought to determine if differential sensitivity may become apparent over a broader range of drug exposure. To this end, the viability of control and RB-depleted cells was assessed following 96-hour exposure to a concentration range of 0.01-100μM of each PARP inhibitor (Fig. 1C). RB-deficient cells exhibited reduced viability compared to control cells following exposure to four inhibitors of PARP (Rucaparib, Veliparib, AG-14361, and AZD2461), but not to the remaining three (INO-1001, 3-Aminobenzamide, and PJ34-HCl). The six inhibitors that selectively reduced viability of RB-depleted cells all target both PARP1 and PARP2, while the three that do not are described to selectively inhibit PARP1 but not PARP2 (Wahlberg et al. 2012; Ali et al. 2016). Interestingly, immunofluorescence-based analysis of γH2AX staining intensity in control and RB-depleted cells verify that the PARP1/2 inhibitor Rucaparib, but not Veliparib, selectively promotes DNA damage in RB-deficient cells (Supplemental Fig. S1C,D). A key distinction between Veliparib and Rucaparib is the mechanism of action by which these drugs inhibit PARPs: Rucaparib, like Olaparib and Talazoparib, perturb PARP function by trapping it on DNA (Zandarashvili et al. 2020). In contrast, Veliparib is an enzymatic inhibitor of PARP function that does not trap the enzyme on DNA (Huang and Kraus 2022). Together these data demonstrate that RB-deficient cells are generally sensitive to combined inhibition of PARP1 and PARP2. These data additionally raise the possibility that the increased DNA damage seen in RB-deficient cells may stem from lesions caused not merely by loss of PARP1 and PARP2 function, but from their sustained association with DNA when inhibited.

To validate PARP inhibition as an approach to specifically sensitize RB-deficient cells to high levels of DNA damage, we next measured γH2AX foci formation in a CRISPR-engineered RB1 knockout cell line ((Nicolay et al. 2015); RPE RB^KO^) with and without RB re-introduction via an inducible Halo-tagged RB construct (RB-Halo; Fig. 2A,B; Supplemental Fig. S2A,B). Cells were exposed to PARP inhibitors Olaparib, Rucaparib, Talazoparib, or Veliparib for 48h and analyzed for γH2AX foci. Here, γH2AX-positive DNA damage foci were quantified per nuclei and cells exhibiting greater than 5 foci were considered to have enhanced DNA damage. Similar to results from shRNA-mediated depletion of RB, RPE RB^KO^ cells exhibit a differential increase in DNA damage following inhibition of PARP via Olaparib, Rucaparib or Talazoparib, but not Veliparib (Fig. 2C,D; Supplemental Fig. S2C). Critically, re-introduction of RB via ectopic expression of Halo-tagged RB construct (RB-Halo) is sufficient to rescue DNA damage in RPE RB^KO^ cells exposed to Olaparib, indicating that sensitivity to PARP inhibition is dependent on loss of RB (Fig. 2E,F).

**Figure 2.**
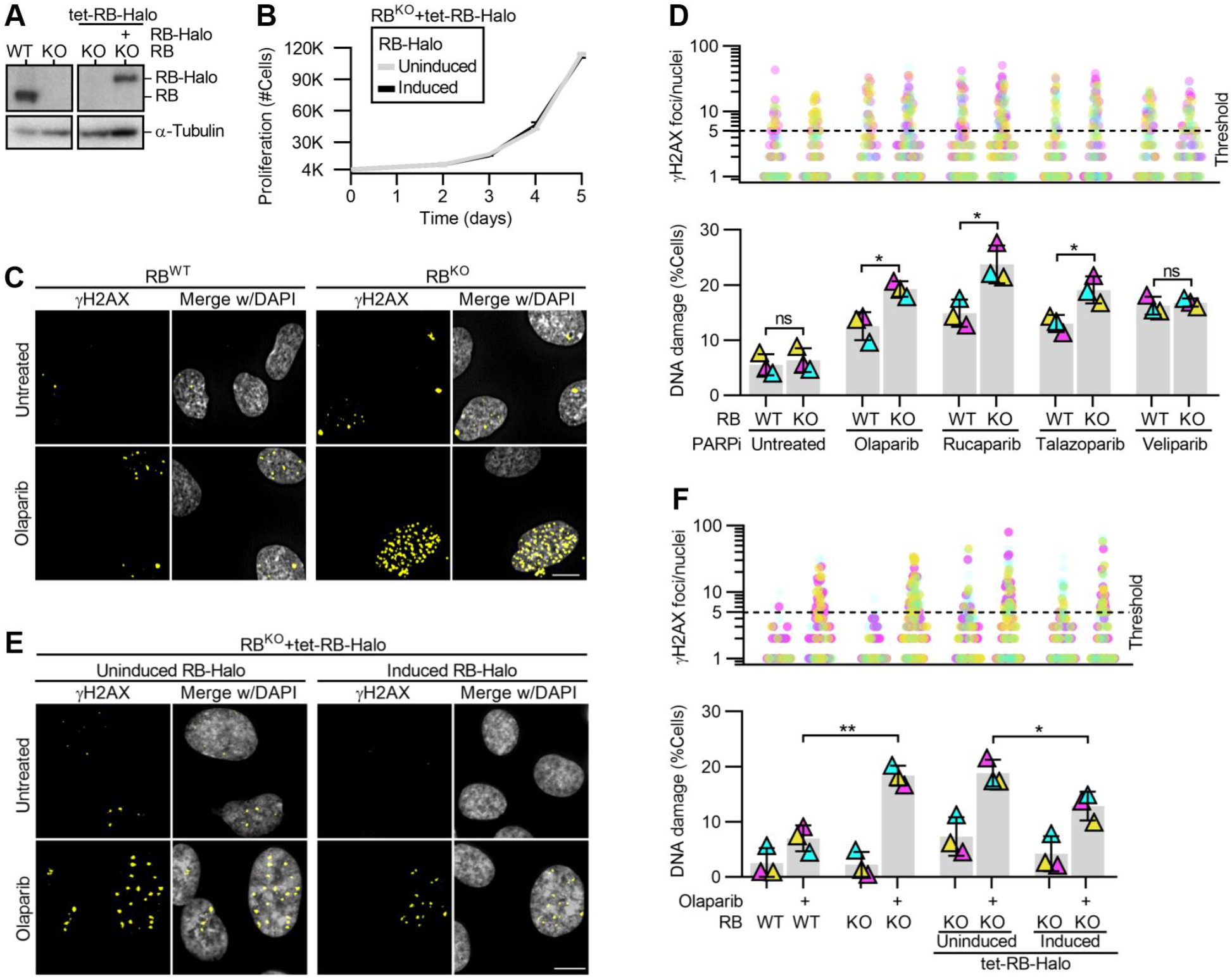
PARP inhibition sensitizes RB-deficient cells to high levels of DNA damage. *(A)* Western blot analysis of control (WT) and RB-null (KO) RPE cells with and without doxycycline inducible RB-Halo (tet-RB-Halo). Cells were induced to express RB-Halo, as indicated (+). *(B)* Quantification of cell number in proliferative populations of RPE RB^KO^ tet-RB-Halo cell number with and without doxycycline induction. *(C, D)* Representative images and quantification of γH2AX foci in RPE RB^WT^ and RB^KO^ cells with and without 48h incubation with the indicated PARP inhibitors. *(E, F)* Representative images and quantification of γH2AX foci in RPE RB^KO^ tet-RB-Halo cells with and without doxycycline induced RB-halo expression and following 48h incubation with Olaparib. *(D, F)* show number of γH2AX foci per cell (top) and percent of cells with ≥5 damage foci (bottom). Scale bars are 10μm. Experiments were performed and statistics calculated between biological triplicates. Error bars represent standard deviation between replicates. (*) p < 0.05; (**) p < 0.01; (ns) non-significant p > 0.05.

### Accumulation of DNA damage following RB loss and PARP inhibition is replication dependent

RB-deficient cells exhibit slow or stalled replication fork progression (Bester et al. 2011; Manning et al. 2014). The PARP enzymes respond to replication stress by placing poly ADP ribose (PAR) modifications to initiate the DNA damage response (Ame et al. 2004) Additionally, PARylation of substrates is necessary for efficient fork restart and Okazaki fragment processing during replication (Sugimura et al. 2008; Bryant et al. 2009; Hanzlikova et al. 2018; Hanzlikova and Caldecott 2019; Vaitsiankova et al. 2022). These reports suggest that RB-deficient cells may rely on PARP-catalyzed PARylation to process their replication stress. To test this hypothesis, we examined cells in S-phase for evidence of PARylation. Control and RB-depleted cells were pulse-labeled with EdU for 30min to enable identification of actively replicating cells, then fixed and immunostained for PAR. EdU was detected via Click-chemistry conjugation of a fluorophore, and DNA detected with DAPI. Intensity of PAR staining per nuclei was measured and compared between actively replicating cells in the control and RB-depleted populations. We find that S-phase cells exhibit an increase in nuclear PAR levels following RB depletion (Fig. 3A,B). Similar results are seen if cells are treated with the PARG inhibitor PDD 00017273 (PARGi), to prevent turnover of PARylation marks (Hanzlikova et al. 2018; Vaitsiankova et al. 2022). Critically, inhibition of replication with emetine (Lukac et al. 2022), prevents the accumulation of PAR in RB-depleted S-phase cells (Fig. 3A,B). These data indicate that during replication, RB-deficient cells experience stress that is sufficient to induce a PARP-dependent response.

**Figure 3.**
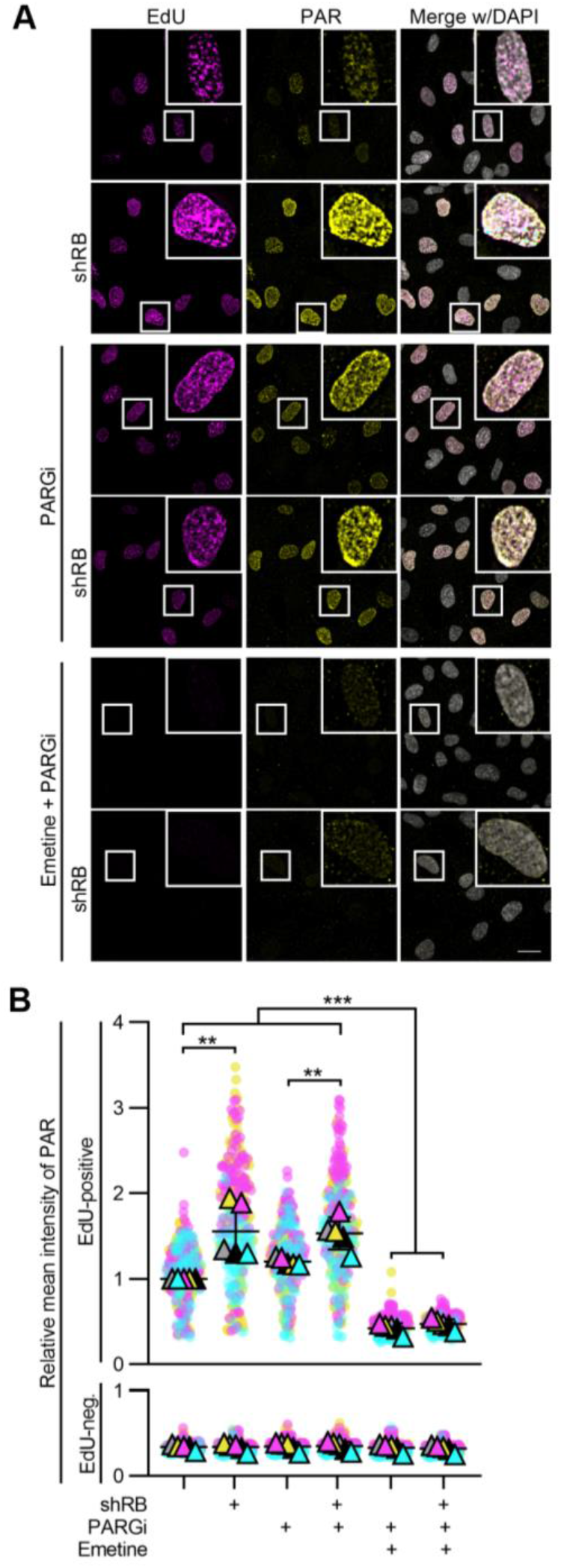
RB loss promotes replication dependent PARylation. *(A, B)* Representative images and quantification of EdU and PAR intensity per cell for RPE tet-shRB cells with and without doxycycline induction of shRB (shRB). Cells were untreated or treated with 2mM Emetine for 1h, and/or 10μM PARG inhibitor for 30min. All conditions were incubated with EdU for the final 30 minutes to label actively replicating cells. Scale bar is 20μm. PAR intensity data are normalized to the EdU-positive, untreated RPE cells. Error bars represent standard deviation between five biological replicates. Statistical analysis was performed between all biological replicates. (**) p < 0.01; (***) p < 0.001.

If replication stress in RB-deficient cells underlies their sensitivity to PARP inhibitors, sites of DNA damage that follow PARP inhibition should correspond with sites of replication stress. To test this possibility, we monitored both replication stress and accumulation of DNA damage concurrently. Replication protein A (RPA) binds to single-stranded DNA, and is phosphorylated at serine-33 (pRPA) in response to replication stress and replication-associated DNA damage (Marechal and Zou 2015). Following 48h exposure to PARP inhibitors Olaparib, Rucaparib or Talazoparib, we found a 2- to 6-fold increase in the fraction of RB-depleted cells with 5 or more pRPA foci, compared to controls (Supplemental Fig. S3A,B; Fig. 4A,B). To confirm these results were specific to RB loss and not the result of potential off-target effects of the RB-targeting shRNA, we also analyzed pRPA and DNA damage foci following siRNA-mediated depletion of RB (Supplemental Fig. S4B-F) and in RB-null Osteosarcoma cells (Supplemental Fig. S4G-J). In both systems, we find results comparable to that described for shRNA-mediated depletion of RB: that RB loss sensitizes cells to high levels of both replication stress and DNA damage following PARP1/2 inhibition. Notably, pRPA foci in RB-deficient, PARP1/2-inhibited cells frequently co-localize with γH2AX foci (Supplemental Fig. S4A,F,J), supporting a model whereby DNA damage is a consequence of unresolved replication stress.

**Figure 4.**
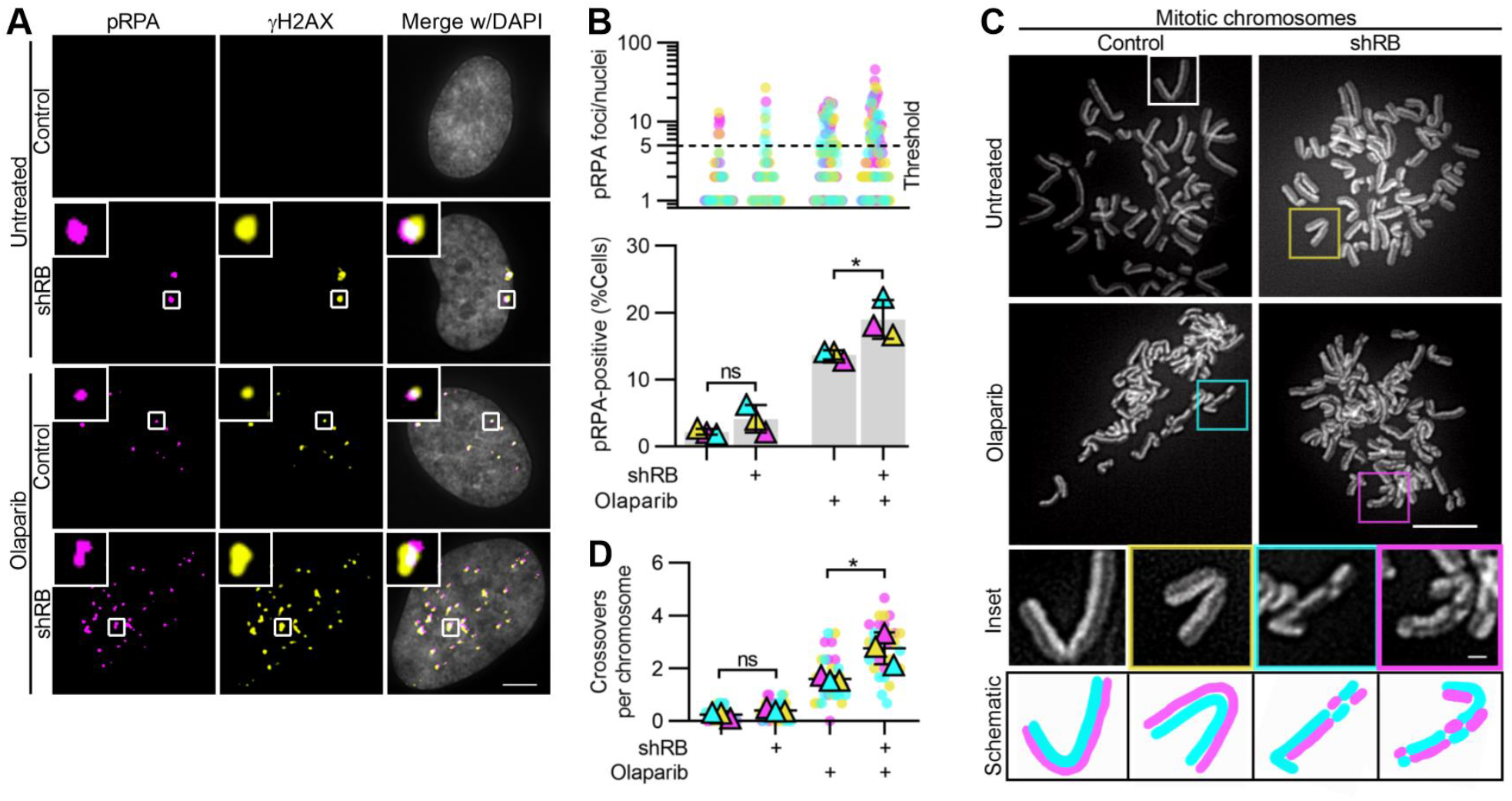
DNA damage accumulates at sites of replication stress following PARP inhibition and RB depletion. *(A, B)* Representative images and quantification of pRPA and γH2AX foci in RPE tet-shRB cells with (shRB) or without (control) doxycycline induction of shRB, following 48h incubation with Olaparib, as indicated. Scale bar is 5μm. *(B)* shows number of pRPA foci per cell (top) and percent of cells with ≥5 damage foci (bottom). Experiments were performed and statistics calculated between biological triplicates. Error bars represent standard deviation between biological replicates. *(C, D)* Representative images and quantification of mitotic crossover events in RPE tet-shRB cells stained with acridine orange. Cells were cultured in 15uM BrdU for 48h and incubated with Olaparib, as indicated, for the final 24h before fixation. Scale bar is 10μm. Insets represents 4x enlargements of a single chromosome, with 1μm scale bar. Diagrams illustrate crossover events present in the inset. Experiments were performed in biological triplicate. Statistics were performed between the average number of crossover events per replicate (*) p < 0.05; (ns) non-significant p > 0.05.

To further define the extent to which RB loss sensitizes cells to PARP inhibition, we performed a sister chromatid exchange assay, differentially staining sister chromatids with acridine orange. In this assay chromosome arm crossovers are a readout of homologous recombination-dependent repair of DNA breaks such that the number of crossovers indicate the frequency of DNA double strand breaks in the preceding S/G2 phases of the cell cycle. We found that RB-depleted cells treated with the PARP inhibitor Olaparib for 24 hours display a significant increase in the number of crossovers per chromosome compared to controls (Fig. 4C,D). Together these data indicate that cells lacking RB are sensitive to the acquisition of replication-dependent DNA double strand breaks following PARP inhibition.

### Persistent replication in the presence of damage perpetuates genomic instability and compromises cell viability

To examine the consequence of extensive DNA damage and investigate the possibility that cell cycle defects correspond with continued replication or translesion repair, cells were briefly pulsed with EdU, followed by examination of mitotic cells. RPE cells spend 2 to 4 hours in G2 following completion of replication before mitotic entry. As a result, cells that enter mitosis during a two-hour EdU pulse are not expected to incorporate EdU. Consistent with this, only ~5% of control mitotic cells exhibit EdU foci following this short pulse. In contrast, ~25% of RB-depleted, PARP1/2-inhibited cells display one or more EdU foci, indicative of under-replicated DNA. Incompletely replicated DNA is susceptible to breaks when chromatin compacts in preparation for mitosis (Lezaja and Altmeyer 2021). Indeed, we found that EdU foci in mitotic RB-depleted and PARP1/2 inhibited cells frequently co-localize with γH2AX (Fig. 5A,B). Consistent with the presence of chromatin breaks during mitosis, interphase RPE RB^KO^ cells treated with PARP inhibitors Olaparib, Rucaparib or Talazoparib for 48 hours exhibit a high frequency of micronuclei. RB loss alone has previously been shown to lead to whole chromosome segregation errors (Manning and Dyson 2011). However, the majority of the micronuclei that result from combined loss of RB and PARP1/2 function lack centromeres, indicating that the increase in micronuclei result from chromatin fragments that fail to incorporate into the main nucleus following mitotic exit, and not from an increase in whole chromosome segregation errors (Fig. 5C,D).

**Figure 5.**
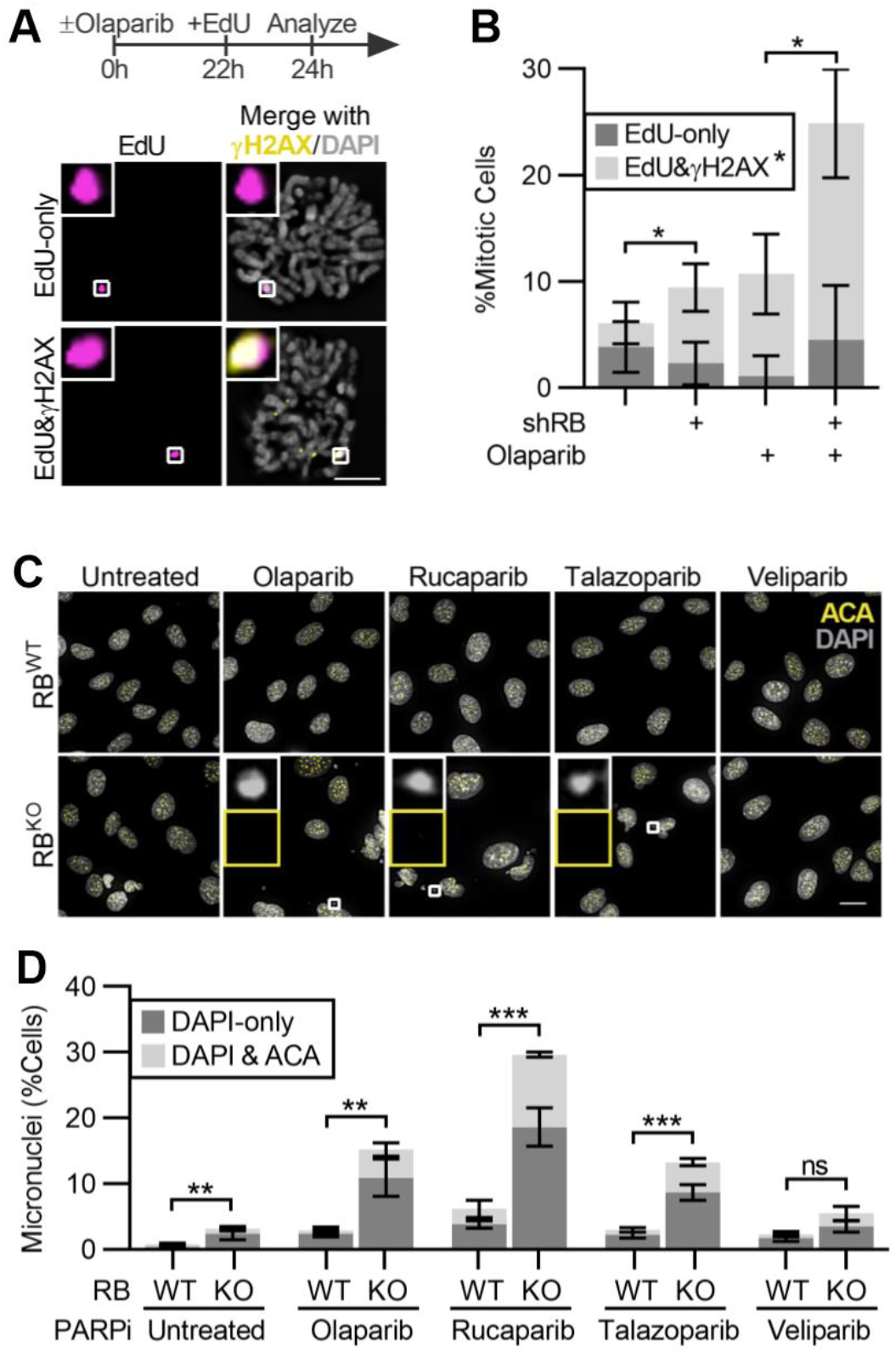
Continued proliferation of RB-deficient cells in the presence of PARP inhibitor promotes genome instability. *(A, B)* Representative images and quantification of EdU and γH2AX foci in mitotic RPE tet-shRB cells with (shRB) or without (control) doxycycline induction of shRB, incubated with 2.5μM Olaparib for 24h, and cultured in 10μM EdU for 2h before fixation. Images and analysis reflect analysis of mitotic cells. Scale bar is 10μm. Experiment was performed in triplicate. Error bars represent standard deviation between biological replicates. Statistical analysis was performed for co-localized γH2AX and EdU foci, across replicates. *(C, D)* Representative images and quantification of the fraction of control (RB^WT^) and RB-null (RB^KO^) cells with micronuclei following incubation with the indicated PARP inhibitors for 48h. Scale bar is 20μm. Experiment was performed in triplicate. Error bars represent standard deviation between biological replicates and statistical analysis is performed between replicates. (*) p < 0.05; (**) p < 0.01; (***) p < 0.001; (ns) non-significant p > 0.05.

Micronuclei are not only a consequence of genome instability but can serve to perpetuate further genomic lesions (Crasta et al. 2012; Soto et al. 2018). Therefore, to assess the long-term impact of increased genomic instability on RB-depleted cells, we monitored the replicative capacity of cells with and without PARP1/2 inhibition and find that continued cell cycle progression can not be maintained following both RB loss and PARP1/2 inhibition. When incubated in media supplemented in EdU for 24 hours, nearly 100% of control and RB-depleted cells, and over 80% of cells treated with PARP1/2 inhibitor alone, incorporate EdU, indicating that these populations are highly proliferative. In contrast, following 48 hours of Olaparib treatment, proliferation of RB-deficient cells is significantly compromised, with only ~40% of the cells within the population remaining competent to incorporate EdU (Fig. 6A,B). Consistent with decreased replication, progression to mitosis is similarly reduced in RB-depleted cells in which PARP1/2 has been inhibited (Fig. 6C). To next determine if PARP1/2 inhibition is cytotoxic for RB-deficient cells, or merely cytostatic, we assessed populations of control and RB-depleted cells for cell death following Olaparib treatment. PARP1/2 inhibition impairs caspase-dependent mechanisms of cell death (Zhang et al. 2012; Tsikarishvili et al. 2021) making readouts of apoptosis ineffective for this system. We therefore opted for live cell imaging to assess the frequency at which cells exhibit blebbing and loss of anchorage, indicative of cell death. Consistent with viability assays (Fig. 1C), live cell imaging confirms that concentrations of PARP1/2 inhibitor that are sublethal for control cells, increase death of RB-depleted and RPE RB^KO^ cells within 72h (Fig. 6D-F). Similar results are seen for RB-null SAOS2 Osteosarcoma cells where dead/dying cells are not apparent in control population but become prevalent following PARP1/2 inhibition (Fig. 6G,H).

**Figure 6.**
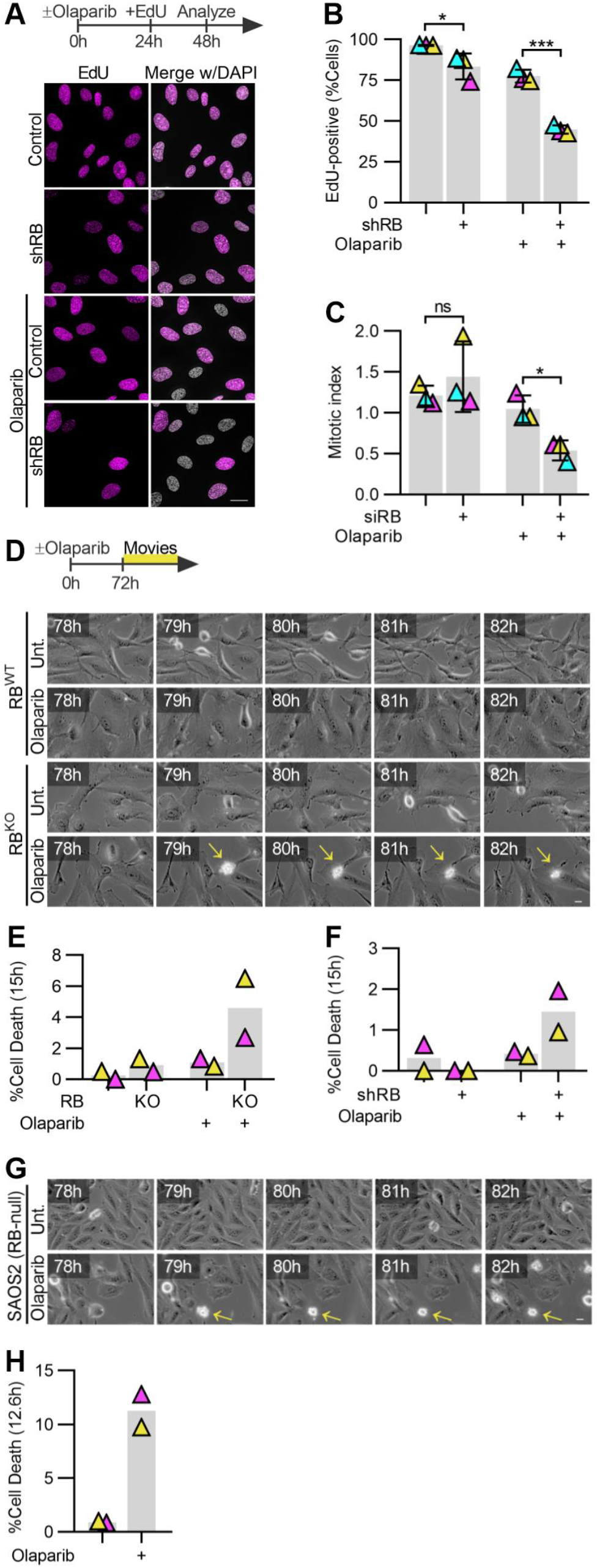
PARP inhibition synergizes with RB loss to compromise cell viability. *(A, B)* Representative images and quantification of replication-competent in RPE tet-shRB cells with (shRB) or without (control) doxycycline induction of shRB, following incubation with 2.5μM Olaparib for 48h. Cells were incubated with 10μM EdU for the final 24h before fixation. Scale bar is 20μm. *(C)* Quantification of the fraction of RPE cell populations in mitosis following siRNA-based depletion of RB, then incubation in 2.5μM Olaparib for 48h. Experiments were performed in triplicate and error bars represent the standard deviation between biological replicates. Statistical analysis was performed between replicates. *(D, E)* Representative still frames and quantification of cell death from live cell imaging of control (RB^WT^) and RB-null (RB^KO^) cells, *(F)* in RPE tet-shRB cells with (shRB) or without (control) doxycycline induction of shRB, and *(G)* RB-null SAOS2 osteosarcoma cells incubated with or without Olaparib. Scale bars are 20μm. Timestamps indicate time since PARP inhibitor addition. Live cell imaging was performed in biological duplicate. (*) p < 0.05; (***) p < 0.001; (ns) non-significant p > 0.05.

## Discussion

Together, this study defines an increased sensitivity of RB-deficient cells to inhibition of PARP1/2 enzymes and mechanistically links this phenotype with replication-induced DNA damage. We show that RB-deficient cells experience stress during replication that promotes robust and sustained PARylation. Following PARP1/2 inhibition, DNA lesions that would otherwise slow replication in a PAR-dependent manner appear to be bypassed as cells progress through S phase. Using a combination of acute depletion of RB and constitutive RB knockout, we show that when PARP1/2 is inhibited sites of replication stress accumulate DNA damage during S phase and that persistent or incomplete replication during mitotic chromatin compaction sensitizes G2/M cells to further DNA damage. The resulting genome instability is evident in fragmented chromatin that accumulate in micronuclei in subsequent cell cycles. In line with previous studies that show RB loss is synthetic lethal with PARP inhibition in osteosarcoma cells (Velez-Cruz et al. 2016; Jiang et al. 2020; Zoumpoulidou et al. 2021), we find that these assaults to genomic integrity correspond with reduced proliferation and increased cell death both in contexts where RB is experimentally depleted, and in cancer cells where the RB1 gene is deleted. These data posit that defects in RB-deficient cells’ capacity to restrain replication in response to stress and/or damage may underlie this synergy. Together with work by (Zoumpoulidou et al. 2021), our data provide evidence that RB status is a clinically relevant biomarker for selection of PARP inhibitors and suggest that RB-deficient cells may be similarly sensitive to additional modulators of replication stress and fork restart (Witkiewicz et al. 2018; Ubhi and Brown 2019).

Of the nine PARP inhibitors evaluated in this study, inhibitors targeting both PARP1 and PARP2, but not those specific for PARP1, had a differential impact on RB-deficient cell viability, suggesting that in the context of this study PARP1 and PARP2 have redundant roles. This is consistent with reports implicating both PARP1 and PARP2 as early sensors of DNA damage and the first line of defense against genomic instability (Hottiger 2015). PARP is activated by binding to sites of single stranded DNA, PARylating various substrates to stabilize replication forks (Yang et al. 2004) and initiate DNA damage repair pathways (Okano et al. 2003). In addition, PARP1 and PARP2 have important roles in regulating Base excision repair (BER, Ronson et al. 2018) pathways and non-homologous end joining (NHEJ)-based repair of DNA double strand breaks (Luijsterburg et al. 2016). This role in NHEJ makes both PARP1 and PARP2 clinically relevant targets in tumors where the complementary double strand break repair pathway, homologous recombination (HR), is already compromised (Bryant et al. 2005; Farmer et al. 2005). Given the previously described role for RB in HR-dependent DNA damage repair pathways (Marshall et al. 2019), it is tempting to speculate that synergy between RB loss and PARP inhibition occurs as a result of dual inhibition of HR and NHEJ. However, this model fails to account for the persistent replication during G2/M that is apparent when both RB and PARP1/2 functions are compromised, but not when either is lost alone. Instead, our findings are consistent with a model whereby PAR-dependent signaling is necessary to sustain fork stability and prevent replication fork progression when DNA lesions are present. In the absence of PAR, RB-deficient cells may bypass such replication impediments, resulting in under-replicated, single stranded regions of DNA that recruit pRPA and are the sites of continued replication late in G2/M. This model is supported by the replication-dependent nature of pRPA and γH2AX accumulation and evidence of underreplicated regions of DNA in mitotic cells in which both RB and PARP function are compromised. This model is also consistent with reports demonstrating that RB loss facilitates translesion synthesis in the presence of crosslinking agents (Bosco et al. 2004) and that RB-null cancers are sensitive to loss of a number of DNA damage repair proteins (Aubry et al. 2020)

### RB loss as a biomarker to predict sensitivity to epigenetic perturbation of replication stress response

Loss or functional inactivation of the retinoblastoma tumor suppressor protein RB is common in a variety of human cancers (Burkhart and Sage 2008; Peifer et al. 2012; George et al. 2015; Sanchez-Vega et al. 2018). Genomics (Burkhart and Sage 2008; George et al. 2015) and transcriptomics (Chen et al. 2019) analysis of cancer patients reveal that RB1-alterations predict poor clinical outcomes, raising the need for targeted therapies. Synthetic lethality is a phenomenon where alterations in one gene hypersensitize cells to alterations in another gene by means of pathway dependence or redundance, and it presents an exciting avenue to target RB-deficient cancers. Recent reports describe synthetic lethal interactions of RB-deficient cells with epigenetic modulators including inhibitors of Aurora kinases (Gong et al. 2019; Oser et al. 2019; Yang et al. 2022), and EZH2 (Ishak et al. 2016). Our data add to a growing body of work showing that RB-deficient cells are exquisitely sensitive to PARP inhibition.

## Materials and Methods

### Cell culture, transfection, and immunofluorescence

hTERT RPE-1 cells and derivatives were cultured in DMEM (GenClone) supplemented with 10% FBS (Sigma) and 1% penicillin/streptomycin (Gibco). RPE RB^KO^ and parental RPE cells were kindly provided by Dr. Nick Dyson (Mass General Research Institute, MGH). SAOS-2 cells were cultured in McCoy’s 5A Medium (Gibco) supplemented with 15% FBS (Sigma) and 1% penicillin/streptomycin (Gibco). All cells were maintained at 37°C and 5% CO_2_. Acute RB depletion was obtained through addition of 2 μg/mL doxycycline to induce expression of an RB-targeting shRNA (Manning and Dyson 2011), or alternatively via reverse-transfection of a pool of four RB-targeting siRNA sequences for 48h, as previously described (Manning et al. 2014). For all experiments, siRNA- or shRNA-driven RB depletion was performed 48h prior to treatment with inhibitors to PARP or other epigenetic modulators. RPE RB^KO^ tet-RB-Halo cells were generated via lentiviral transduction of a pLenti CMV/TO RB-Halo construct. Depletion and/or induced expression of RB was monitored by western blot analysis of whole cell lysates prepared using 2x Laemmli sample buffer (Bio-Rad #1610737) supplemented with 2-mercapto-ethanol (Fisher Scientific #BP176-100, 1:20). Further information on reagents and antibodies used can be found in Supplemental Table S3.

For the epigenetic modulator screen cells were cultured in imaging bottom dishes (Corning #3904) and incubated with individual drugs at a final concentration of 10μM for 48h prior to fixation in 4% PFA and γH2AX immunofluorescence as previously described (Manning et al. 2014). Cells were subsequently counterstained with fluorophore-conjugated secondary antibodies (Invitrogen) and 0.2μg/mL DAPI. 10mg/mL DABCO (Thermo Scientific #112470250) in glycerol-PBS (9:1) antifade reagent was used to stabilize signal. γH2AX and/or pRPA immunofluorescence was performed in cells grown on coverslips and fixed in 4% PFA, as previously described (Manning et al. 2014). Replicating cells were labeled with a pulse of 10μM EdU for 2-24h, as indicated in the text, and visualized using the Click-iT EdU Imaging Kit (InVitrogen #C10637), per the manufacturer’s instructions. To detect PAR, cells were processed as previously described (Vaitsiankova et al. 2022) and incubated with PAR antibody. Cells were counterstained with fluorophore-conjugated secondary antibodies and 0.2μg/mL DAPI, mounted on slides with ProLong Gold, and imaged using a Nikon Eclipse Ti-E microscope equipped with a Zyla sCMOS camera and controlled by NIS Elements software. All experiments were performed in triplicate and a minimum of 30 cells per condition were analyzed. Experimental conditions from within a biological replicate were imaged in parallel and at the same exposure time. Representative images were deconvolved in NIS Elements software.

For live cell imaging, cells were grown on 12-well plates and treated with Olaparib for 72h. Cells were imaged using the Nikon Eclipse Ti-E equipped with perfect focus software and an environmental chamber to maintain 37°C and 5% CO_2_. Phase contrast images were captured every 5 minutes for the final 12-15h of Olaparib incubation. Gamma-adjustments were made for representative movie stills to enable better visualization across time.

### Cell proliferation and viability assays

For each of three biological replicates, cells were incubated with the indicated inhibitor concentrations in technical duplicates. At 96h cells were suspended and manually counted with a hemocytometer. Alternatively, PrestoBlue™ Cell Viability Reagent (InVitrogen #A13261, 1:10 dilution) was added at indicated timepoints and incubated for 3h. Fluorescence was analyzed at 560/590nm ex/em with a PerkinElmer Victor3 1420 plate reader.

### Sister chromatid exchange assay

To differentially label replicated sister chromosomes, cells were incubated in 15μM BrdU for 48h (~2 cell cycles). When indicated, Olaparib was added for the final 24h of BrdU labeling. Chromosome spreads were prepared by treating cells with 0.1μg/mL Nocodazole for 30min to depolymerize microtubules, followed by incubation in 75mM KCl for 16 minutes. Cells were fixed in methanol-acetic acid (3:1) for 20 minutes at 4°C. Fixed cells were then dropped onto slides, dried overnight in the dark, stained with 100μg/mL acridine orange (Molecular Probes #A3568), and mounted in 0.1M Na_2_HPO_4_ and 0.1M KH_2_PO_4_.

### Image analysis and statistics

For the screen, extended depth of focus 2D-projections were generated using NIS Elements software. Nuclear regions were identified based on a DAPI threshold and γH2AX staining intensities measured. Cells were considered damaged if the sum intensity of a given cell was 2-fold or greater than the average intensity observed in the untreated controls. For imaging of cells on coverslips, a Cell Profiler (Stirling et al. 2021) pipeline was generated to first identify nuclei and then detect individual γH2AX-positive foci. Cells were considered damaged if the γH2AX foci count per nuclei was 5 or greater. Thresholds for nuclear EdU staining intensities were set on a per-replicate basis and kept consistent across conditions within the replicate. SuperPlots represent 100 cells scored per experimental condition and triangles indicate averages per biological replicate. Statistical analysis was performed for biological triplicates. Where relevant, technical replicates were averaged before comparisons were made between biological triplicates.

Unless stated otherwise, statistical analyses are two-tailed unpaired student’s t-tests, with P-values indicated in figure legends. For the screen statistical analysis, robust-Z-scores were calculated as previously described (Chung et al. 2008) via the following equation:

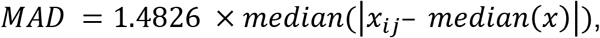

where x_ij_ indicates the fold change of percent damaged cells between RB-depleted and control cells for a given experimental condition (shown in Supplemental Table S1), and x indicates the median among the 79 epigenetic modulators tested. Robust Z-scores are defined by x_ij_ divided by MAD. Hit discovery was assessed with a threshold of Z≥3.

## Acknowledgements

We thank members of the Manning and Flynn Labs for technical assistance and critical reading of the manuscript. This work was supported by a National Institute of Health award R01CA214880 to RLF and American Cancer Society RSG-21-066-01-CCG and Smith Family Awards to ALM.

## Author Contributions

ALM and RLF conceived of the study and designed the experiments. LGZ, MMP, NMH, and HG designed, performed, and analyzed the experiments. ALM and LGZ wrote the manuscript.

