## Supplemental Material for "RB loss sensitizes cells to replication-associated DNA damage by PARP inhibition"

### Supplemental Table S1. DNA damage parameters of screened epigenetic modulators

| Target | Drug | DNA damage (Fold-change relative to untreated control) | | Robust Z-score |
|---|---|---|---|---|
| | | Control | shRB | |
| AURK | CYC116 | 0.11 | 1.64 | 20.04 |
| AURK | JNJ-7706621 | 2.01 | 4.52 | 3.12 |
| AURK | PHA-680632 | 1.31 | 2.70 | 2.87 |
| AURK | VX-680 (Tozasertib) | 3.39 | 3.83 | 1.56 |
| AURK | MK-5108 (VX-689) | 1.52 | 1.69 | 1.54 |
| AURK | CCT129202 | 2.00 | 2.22 | 1.53 |
| AURK | Danusertib (PHA-739358) | 0.96 | 0.96 | 1.38 |
| AURK | MLN8054 | 1.82 | 1.78 | 1.35 |
| AURK | Barasertib (AZD1152-HQPA) | 2.51 | 2.14 | 1.18 |
| AURK | Aurora A Inhibitor I | 2.45 | 1.80 | 1.02 |
| AURK | ZM 447439 | 1.87 | 1.35 | 1.00 |
| AURK | Alisertib (MLN8237) | 2.20 | 1.40 | 0.88 |
| AURK | SNS-314 Mesylate | 2.29 | 1.21 | 0.73 |
| AURK | AMG-900 | 3.69 | 1.85 | 0.70 |
| BRD4 | PFI-1 (PF-6405761) | 1.47 | 4.81 | 4.52 |
| BRD4 | Bromosporine | 0.85 | 2.02 | 3.29 |
| BRD4 | RVX-208 | 0.68 | 1.15 | 2.35 |
| BRD4 | CPI-203 | 0.74 | 1.08 | 2.02 |
| BRD4 | UNC1215 | 0.36 | 0.45 | 1.72 |
| BRD4 | I-BET151 (GSK1210151A) | 1.67 | 1.69 | 1.40 |
| BRD4 | I-BET-762 | 1.33 | 1.30 | 1.35 |
| BRD4 | SGC-CBP30 | 0.69 | 0.64 | 1.28 |
| BRD4 | OTX015 | 2.94 | 1.65 | 0.78 |
| BRD4 | (+)-JQ1 | 1.87 | 0.84 | 0.62 |
| DNMT | Zebularine | 0.57 | 2.25 | 5.45 |
| DNMT | Azacitidine | 0.19 | 0.70 | 5.24 |
| DNMT | SGI-1027 | 0.43 | 1.36 | 4.36 |
| DNMT | RG108 | 0.49 | 0.68 | 1.94 |
| DNMT | Lomeguatrib | 1.30 | 0.80 | 0.86 |
| EZH2 | 3-Deazaneplanocin A (DZNeP) | 0.36 | 1.78 | 6.88 |
| EZH2 | EPZ-6438 (Tazemetostat) | 0.34 | 1.15 | 4.66 |
| EZH2 | EPZ5676 | 0.48 | 1.18 | 3.39 |
| EZH2 | Entacapone | 0.84 | 1.33 | 2.20 |
| EZH2 | MM-102 | 0.67 | 1.01 | 2.07 |
| EZH2 | EPZ004777 | 0.67 | 0.76 | 1.56 |
| EZH2 | SGC 0946 | 0.47 | 0.50 | 1.49 |
| HDAC | RGFP966 | 0.92 | 2.89 | 4.36 |
| HDAC | Rocilinostat (ACY-1215) | 1.35 | 3.58 | 3.66 |
| HDAC | Resminostat | 1.06 | 2.71 | 3.53 |
| HDAC | Givinostat (ITF2357) | 3.75 | 5.36 | 1.98 |
| HDAC | Scriptaid | 1.97 | 2.19 | 1.54 |
| HDAC | Belinostat (PXD101) | 5.12 | 5.08 | 1.37 |
| HDAC | PCI-24781 (Abexinostat) | 1.76 | 1.66 | 1.31 |
| HDAC | Trichostatin A (TSA) | 7.47 | 6.17 | 1.14 |
| HDAC | Panobinostat (LBH589) | 10.72 | 6.17 | 0.80 |
| HDAC | TMP269 | 1.08 | 0.59 | 0.75 |
| HDAC | Entinostat (MS-275) | 5.53 | 2.64 | 0.66 |
| HDM | GSK J4 HCl | 1.65 | 6.44 | 5.40 |
| HDM | IOX1 | 0.53 | 1.09 | 2.86 |
| HDM | Tranylcypromine (2-PCPA) HCl | 1.05 | 1.29 | 1.70 |
| HDM | OG-L002 | 0.91 | 1.02 | 1.56 |
| JAK | AZD1480 | 0.51 | 2.52 | 6.91 |
| JAK | Ruxolitinib (INCB018424) | 0.34 | 1.09 | 4.42 |
| JAK | Gandotinib (LY2784544) | 2.49 | 5.67 | 3.15 |
| JAK | Tofacitinib (CP-690550) | 1.19 | 1.93 | 2.24 |
| JAK | CEP-33779 | 0.46 | 0.31 | 0.92 |

| | | | | |
|---|---|---|---|---|
| JAK | CYT387 | 2.92 | 1.33 | 0.63 |
| PARP1/2 | Olaparib (AZD2281) | 1.14 | 3.10 | 3.76 |
| PARP1/2 | Talazoparib (BMN 673) | 4.64 | 11.24 | 3.35 |
| PARP | Rucaparib (AG-014699) | 1.39 | 2.61 | 2.60 |
| PARP1 | AG-14361 | 0.97 | 1.71 | 2.44 |
| PARP | PJ34 HCl | 0.45 | 0.70 | 2.14 |
| PARP1/2 | Veliparib (ABT-888) | 1.74 | 2.43 | 1.93 |
| PARP | 3-Aminobenzamide | 0.92 | 0.88 | 1.32 |
| PARP | AZD2461 | 0.72 | 0.64 | 1.24 |
| PARP | INO-1001 | 0.47 | 0.36 | 1.07 |
| Other | SMI-4a | 0.69 | 2.09 | 4.21 |
| Other | C646 | 0.36 | 0.64 | 2.49 |
| Other | Quercetin | 1.34 | 2.36 | 2.44 |
| Other | AZD1208 | 0.46 | 0.80 | 2.41 |
| Other | Resveratrol | 0.90 | 0.99 | 1.53 |
| Other | Sirtinol | 0.98 | 1.07 | 1.50 |
| Other | FG-4592 | 1.02 | 1.01 | 1.37 |
| Other | EX 527 (Selisistat) | 1.08 | 1.03 | 1.32 |
| Other | Iniparib (BSI-201) | 1.58 | 1.26 | 1.10 |
| Other | Procainamide HCl | 0.96 | 0.65 | 0.94 |
| Other | IOX2 | 0.86 | 0.56 | 0.90 |
| Other | CX-6258 HCl | 7.10 | 0.00 | 0.00 |

**Supplemental Table S2. Survival $IC_{50}$ values for screen hits and additional PARP inhibitors**

| Target | Drug | Survival $IC_{50}$ (μM) | | $IC_{50}$ ratio (Control/shRB) |
|---|---|---|---|---|
| | | Control | shRB | |
| AURK | CYC116 | 2.57 | 1.78 | 1.44 |
| AURK | JNJ-7706621 | 1.36 | 1.28 | 1.06 |
| BRD4 | PFI-1 (PF-6405761) | 13.52 | 9.89 | 1.37 |
| BRD4 | Bromosporine | 4.33 | 2.65 | 1.63 |
| EZH2 | DZNeP | >100 | >100 | NA |
| EZH2 | Tazemetostat | 0.34 | 0.75 | 0.45 |
| HDAC | RGFP966 | 56.62 | 29.49 | 1.92 |
| HDAC | Rocilinostat | 8.26 | 4.63 | 1.78 |
| HDAC | Resminostat | 4.80 | 2.35 | 2.04 |
| JAK | AZD1480 | 40.88 | 20.75 | 1.97 |
| JAK | Ruxolitinib | 5.52 | 3.25 | 1.70 |
| JAK | Gandotinib | ≳100 | 10.78 | NA |
| PARP1/2 | Olaparib | 4.78 | 1.74 | 2.74 |
| PARP1/2 | Talazoparib | 0.28 | 0.02 | 14.00 |
| PARP | Rucaparib | 3.28 | 0.80 | 4.10 |
| PARP1 | AG-14361 | 24.46 | 5.64 | 4.33 |
| PARP1/2 | Veliparib | 57.24 | 13.07 | 4.38 |
| PARP | 3-Aminobenzamide | >100 | >100 | NA |
| PARP | AZD2461 | 22.15 | 6.00 | 3.69 |
| PARP | INO-1001 | >100 | >100 | NA |
| PARP | PJ34-HCl | 71.41 | 42.21 | 1.69 |

(NA) Not applicable

**Supplemental Table S3. Drugs and antibodies used**

| Reagent | Company | Catalog number | Concentration |
|---|---|---|---|
| Doxycycline | Thermo Fisher | 446061000 | 2µg/mL |
| Epigenetic Modulator Library | Selleck Chemicals | Z113676 | 10µM |
| Olaparib | Selleck Chemicals | S1060 | 2.5µM |
| Rucaparib | Selleck Chemicals | S1098 | 2.5µM |
| Talazoparib | Selleck Chemicals | S7048 | 20nM |
| Veliparib | Selleck Chemicals | S1004 | 10µM |
| PARG inhibitor | Tocris | 5952 | 10µM |
| Emetine | Sigma | E2375 | 2µM |
| Mouse-RB (clone 4H1) | Cell Signaling | 9009L | 1:1000 dil |
| Mouse-GAPDH | Proteintech | 600004-1 | 1:100000 dil |
| Mouse- α-tubulin | Santa Cruz | sc-32293 | 1:1000 dil |
| HRP-labeled anti-mouse | Cytiva | NA931V | 1:5000 dil |
| Rabbit-γH2AX | Cell Signaling | 2577L | 1:1000 dil |
| Mouse-γH2AX | Millipore | 05-636 | 1:500 dil |
| Rabbit-pRPA | Sigma | PLA0310 | 1:500 dil |
| Human-ACA | Antibodies Inc. | 15-234 | 1:500 dil |
| Mouse-PAR | Trevigen | 4335-MC-100 | 1:250 dil |
| Alexa Fluor 488 anti-rabbit | Thermo Fisher | A-11008 | 1:1000 dil |
| Alexa Fluor 488 anti-human | Thermo Fisher | A-11013 | 1:1000 dil |
| Alexa Fluor 546 anti-rabbit | Thermo Fisher | A-10040 | 1:1000 dil |
| Alexa Fluor 546 anti-mouse | Thermo Fisher | A-11003 | 1:1000 dil |
| Alexa Fluor 680 anti-rabbit | Thermo Fisher | A-21076 | 1:1000 dil |
| Alexa Fluor 680 anti-mouse | Thermo Fisher | A-21058 | 1:1000 dil |
| Acridine Orange | Thermo Fisher | A-3568 | 100µg/mL |

**Supplemental Materials and Methods**

**$IC_{50}$ calculations.** Relative $IC_{50}$ calculations for Supplemental Table 2 were performed in GraphPad Prism software, using four-parameter logistic regression with iterative predictions of the following equation:

$$Y = Bottom + (Top - Bottom) / (1 + 10\^((LogIC50 - X) \times HillSlope)),$$

where X is a concentration (µM) tested for a given drug, Y is the respective %Cell survival, Top and Bottom are plateaus in units of %Cell survival, HillSlope indicates the steepness of the curve, and LogIC50 indicates the logarithm in base 10 of the concentration required to bring the curve down to the point halfway between Top and Bottom.

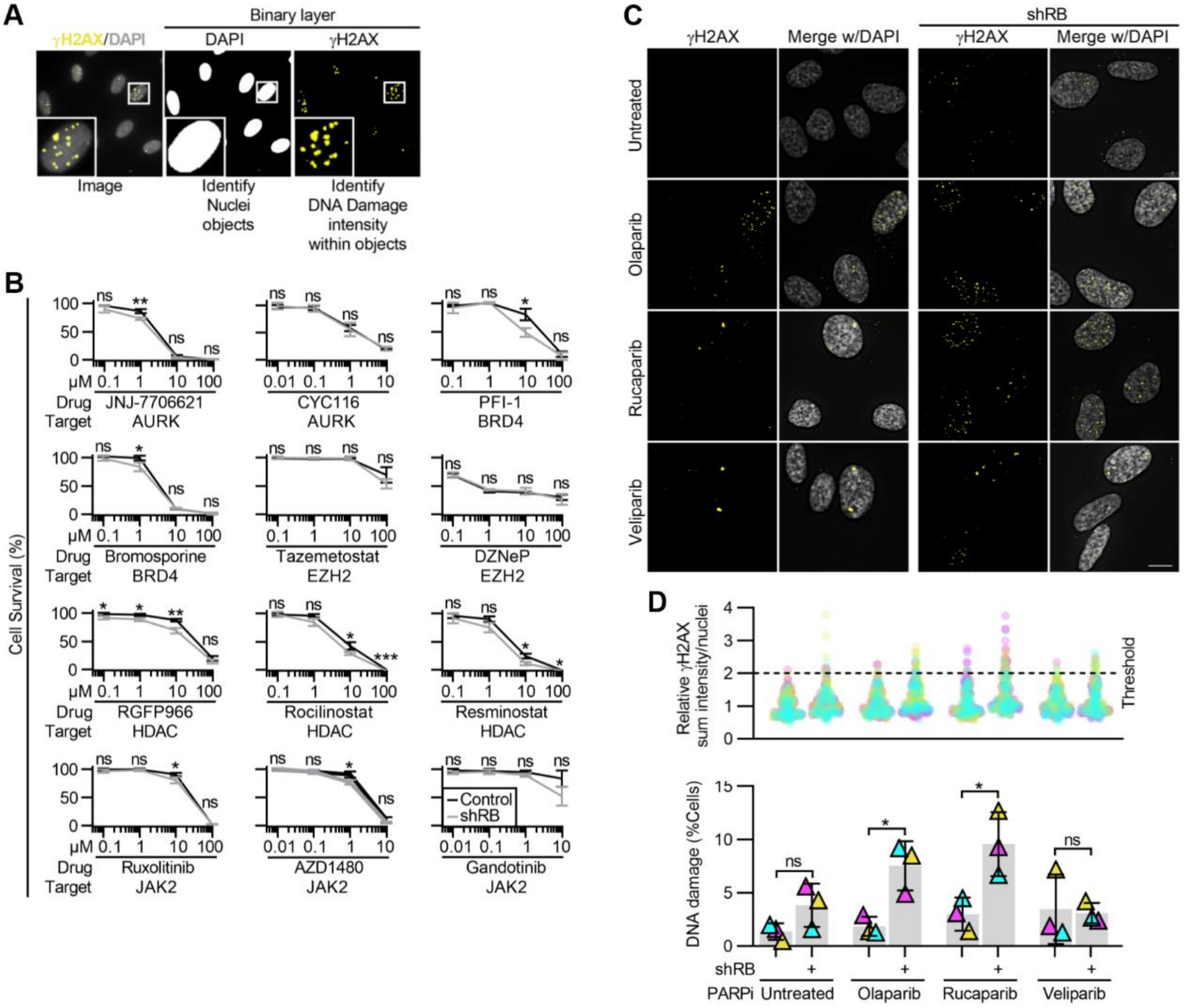


Supplemental Figure 1. Acute depletion of RB renders cells sensitive to a panel of PARP1/2 inhibitors. *(A)* Visualization of NIS Elements analysis indicating nuclei segmentation and DNA damage identification based on γH2AX staining intensity. *(B)* Relative cell survival indicated by metabolic color conversion of PrestoBlue™ reagent, in RPE tet-shRB cells with and without doxycycline induced shRB expression, following incubation with the indicated PARP inhibitors and concentrations. *(C, D)* Representative images and quantification of γH2AX foci in control and shRB cells, following 48h incubation with PARP inhibitors, as indicated. Scale bar is 5μm. *(D)* shows number of γH2AX foci per cell (top) and percent of cells with ≥5 damage foci (bottom). All experiments were performed in biological triplicate with error bars representing standard deviation between replicates, and statistics calculated between biological replicates. (*) $p < 0.05$; (**) $p < 0.01$; (***) $p < 0.001$; (ns) non-significant $p > 0.05$.

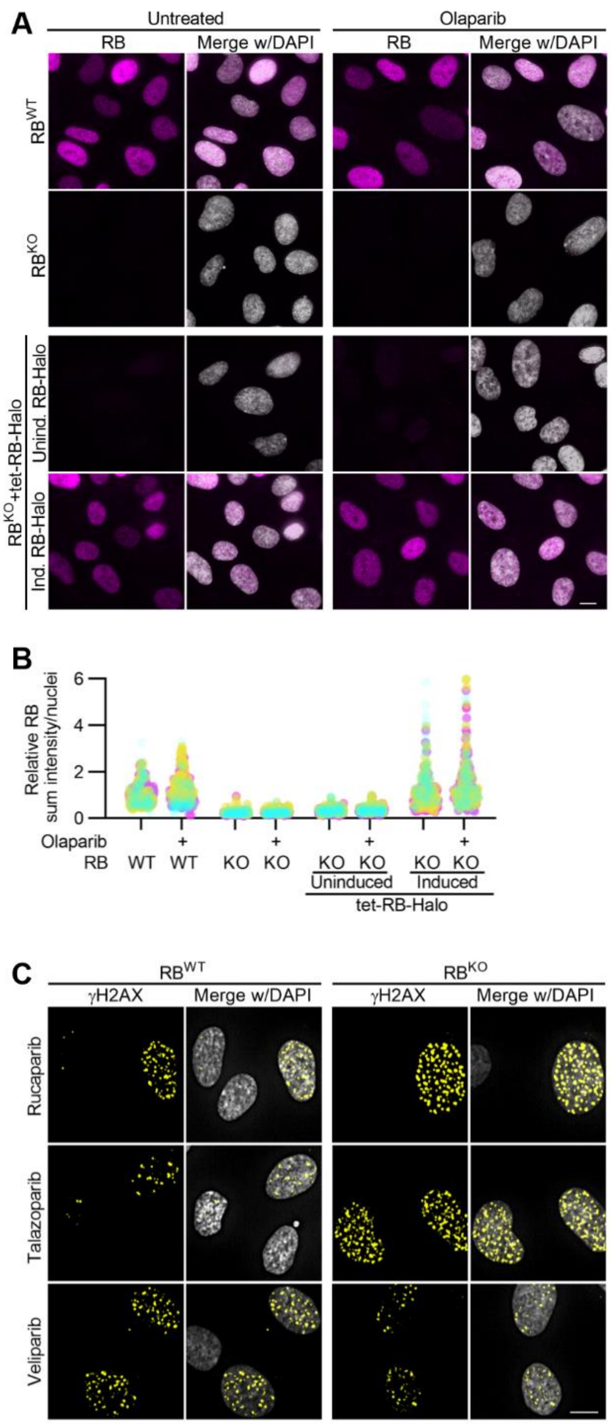


Supplemental Figure 2. Exogenous expression of RB-Halo in RB-null RPE cells rescues PARP sensitivity. *(A, B)* Representative images and quantification of RB levels in control ($RB^{WT}$) and RB-null ($RB^{KO}$) RPE cells with and without doxycycline induced expression of Halo-tagged RB (RB-Halo), following incubation with Olaparib for 48h. *(C)* Representative images of γH2AX foci in $RB^{WT}$ and $RB^{KO}$ RPE cells incubated with PARP inhibitors Rucaparib, Talazoparib or Veliparib for 48h. Quantification is in Fig. 2D. Scale bars are 10μm. All experiments were performed in biological triplicate.

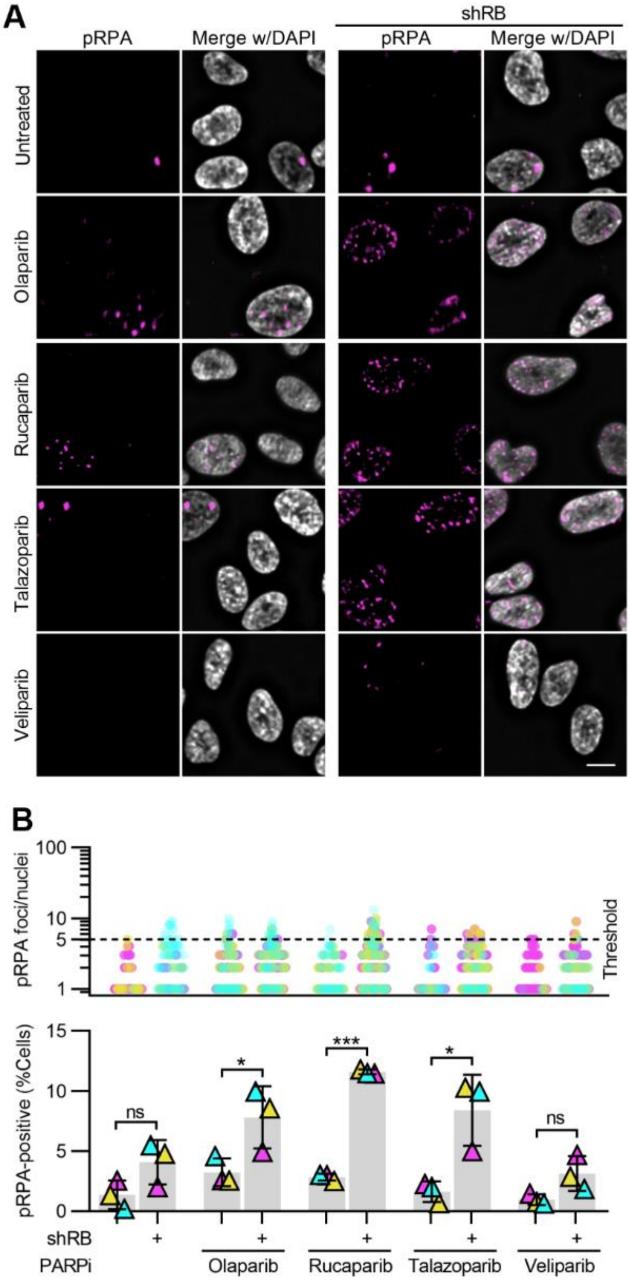


Supplemental Figure 3. RB-depleted cells exhibit replication-dependent DNA damage following PARP inhibition. *(A, B)* Representative images and quantification of pRPA foci in RPE tet-shRB cells with and without doxycycline induced shRB expression, following 48h incubation with PARP inhibitors, as indicated. Scale bar is 10μm. *(B)* shows number of pRPA foci per cell (top) and percent of cells with ≥5 foci (bottom). All experiments were performed in biological triplicate with error bars representing standard deviation between replicates, and statistics calculated between biological replicates. (*) $p < 0.05$; (***) $p < 0.001$; (ns) non-significant $p > 0.05$.

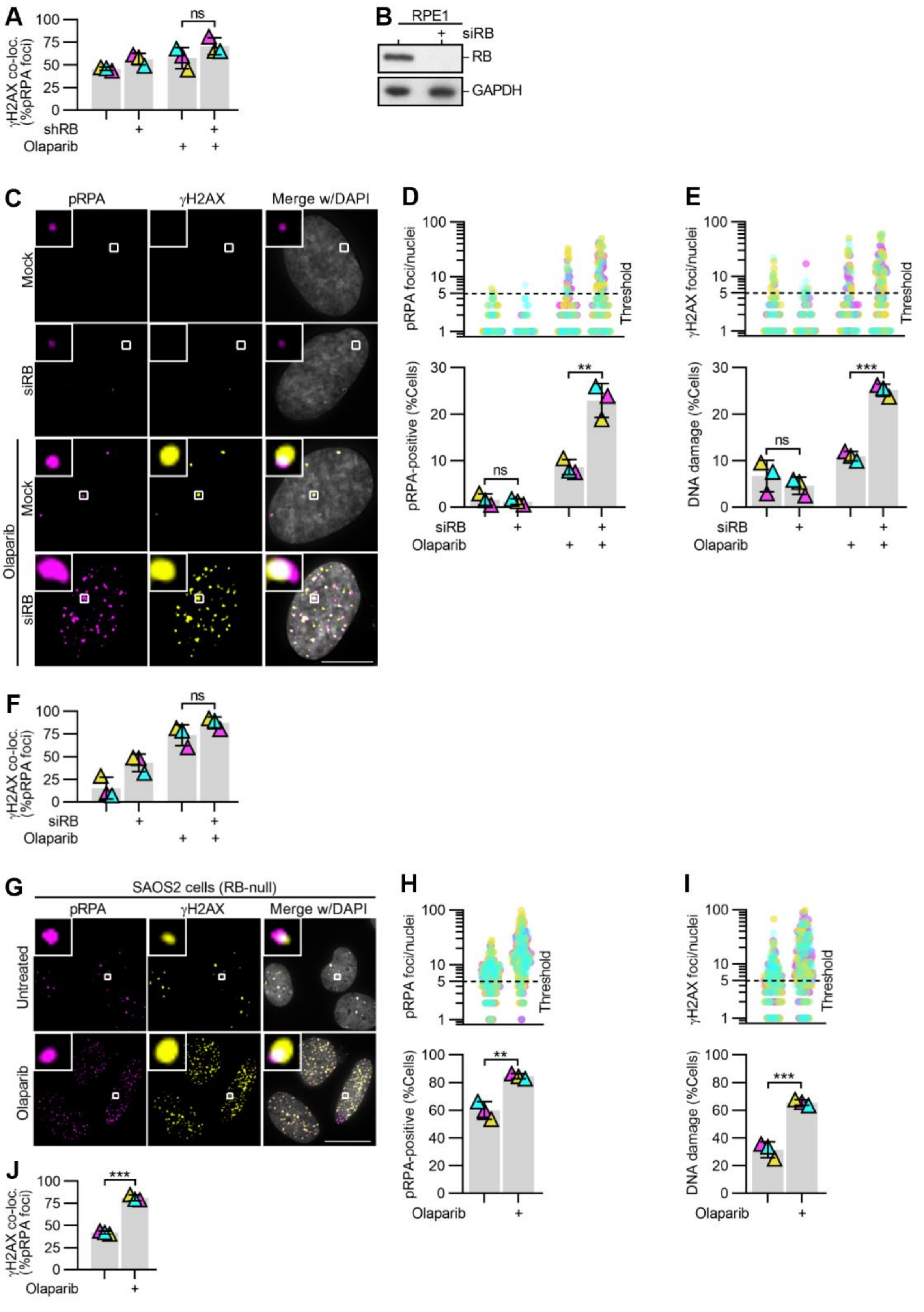


Supplemental Figure 4. PARP inhibition following acute and constitutive loss of RB similarly promote replication-associated DNA damage. *(A)* Quantification of colocalization of γH2AX and pRPA foci for experiments described in Fig. A,B. *(B)* Western blot analysis of RB protein levels following transfection with mock (control) or RB-targeting siRNA pool (siRB). *(C-F)* Representative images and quantification of pRPA and γH2AX foci in mock or siRB transfected RPE cells, following 48h incubation with Olaparib, as indicated. Scale bar is 10μm. *(D, E)* show number of pRPA or γH2AX foci per cell (top) and percent of cells with ≥5 damage foci (bottom). *(F)* Quantification of co-localization of γH2AX and pRPA foci for experiments in C-E. *(G-J)* Representative images and quantification of pRPA and γH2AX foci in RB-null SAOS2 osteosarcoma cells, following 48h incubation with 10μM Olaparib, as indicated. Scale bar is 20μm. *(H, I)* show number of pRPA or γH2AX foci per cell (top) and percent of cells with ≥5 damage foci (bottom). *(J)* Quantification of co-localization of γH2AX and pRPA foci for experiments in G-I. All experiments were performed in biological triplicate with error bars representing standard deviation between replicates, and statistics calculated between biological replicates. (**) $p < 0.01$; (***) $p < 0.001$; (ns) non-significant $p > 0.05$.
